# Changes in Gait Asymmetry May Be Caused by Adaptation of Spinal Reflexes

**DOI:** 10.1101/2024.02.04.578817

**Authors:** Omar Refy, Owen Mo, Jacob Hsu, Douglas J. Weber, Hartmut Geyer

**Author notes:** (co-senior author). Email: dweber2andrew.cmu.edu (co-senior author).

## Abstract

In a recent human study, we found that adaptive changes in step length asymmetry (SLA) are correlated with similar changes in the H-reflex gains of the leg muscles during split-belt treadmill locomotion. While this observation indicated a closer link between gait asymmetry and spinal reflex adaptation, it did not reveal their causal relationship. To better understand this relationship, here we use a neuromuscular model of human walking whose control relies primarily on spinal reflexes. Subjecting the model to split-belt treadmill locomotion with different combinations of belt speed and reflex gain adaptation patterns, we find that belt speed changes increase the variability in SLA but do not result in consistent SLA patterns as observed in human experiments, whereas reflex gain adaptations do. Furthermore, we find that the model produces SLA patterns similar to healthy adults when its reflex gains are adapted in a way similar to the H-reflex changes we observed in our previous human study. The model also predicts SLA patterns similar to the ones observed for cerebellar degeneration patients when the reflexes do not adapt beyond a sudden dip at the time the ipsilateral belt speed is lowered. Our results suggest that SLA does not arise from imposing belt speed changes but requires the adaptation of the reflex gains, and that the dynamic adaptation of these gains may be an essential part of human gait control when encountering unexpected environment changes such as the uneven speed changes in split-belt treadmill locomotion.

## 1 Introduction

The human motor system can adapt gait both permanently and transiently in response to neurologic diseases, injuries, and gait disturbances (Nas, 2015; Li et al., 2018; di Biase et al., 2020). Understanding the neural mechanisms behind this adaptation can not only provide scientists with a deeper knowledge about human gait physiology but also serve clinicians as a guide for correcting gait pathologies when necessary.

Locomotion on a split-belt treadmill with asymmetric belt speeds has become a well established paradigm for studying human gait adaptation in the lab environment (Reisman et al., 2005; Torres-Oviedo and Bastian, 2010). Many of the original works using this paradigm identified key observations about split belt adaptation, such as that it involves sensorimotor calibration (Vazquez et al., 2015) and that supraspinal centers are essential for the adaptation (Dietz et al., 1994; Morton and Bastian, 2006). In more recent years, a number of studies have started to focus on the neural origins of these observations. For instance, Iturralde and Torres-Oviedo (2019) measured electromyography (EMG) from major leg muscle groups during split-belt adaptation and found that, for some muscles, EMG activity is distinctly different in early versus late adaptation. For another example, using electroencephalography (EEG), Jacobsen and Ferris (2023) have found that multiple brain areas are active during split-belt locomotor adaptation. While these and other examples provide evidence that supraspinal centers are involved in gait adaptation, it remains unclear whether and how this involvement affects locomotor control circuits in the spinal cord.

In a recent study, we combined split belt locomotion with H-reflex measurements to study spinal reflexes during gait adaptation in able-bodied humans(Refy et al., 2023). We found that changes in step length asymmetry (SLA) are correlated with similar changes in H-reflex gains of the leg extensor muscles (Fig. 1), suggesting that SLA is accompanied by reflex adaptations. However, these results did not reveal if the adaptations are caused by SLA or the other way around. Some support for the latter case comes from the work of Thompson et al. (2013); Thompson and Wolpaw (2014), who showed that conditioning of reflexes over months can correct gait asymmetry in people with paraplegia due to spinal cord injury. Yet the reflex adaptations we observed occurred over the course of mere minutes and in near lockstep with the SLA changes (Fig. 1).

**Figure 1.**
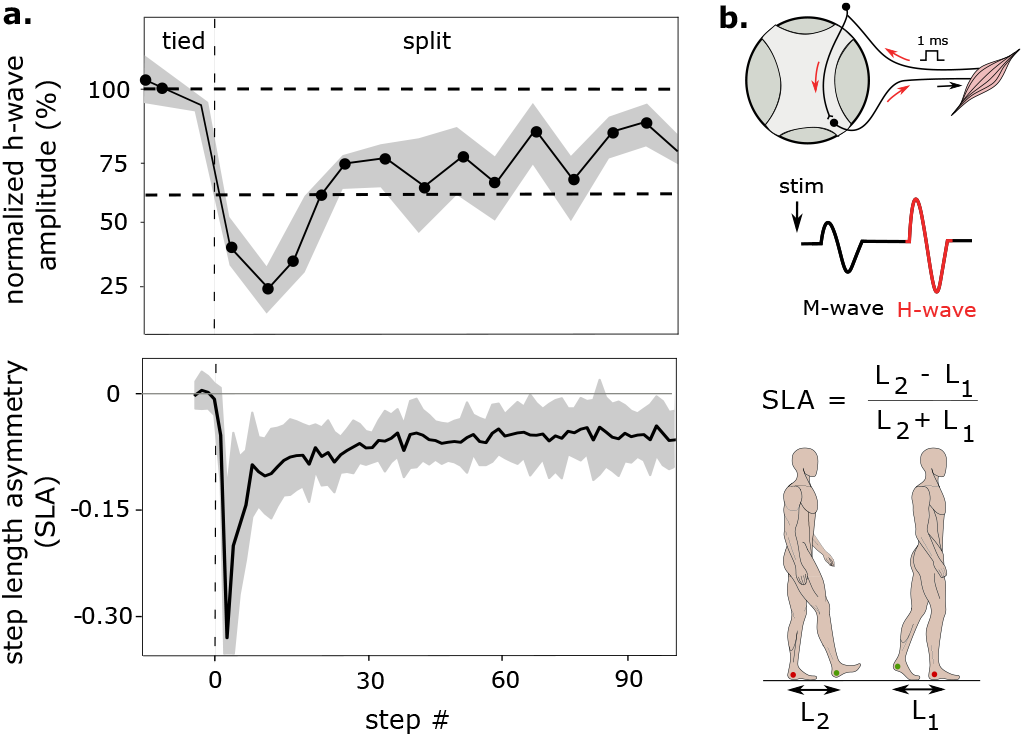
Step length asymmetry (SLA) and H-reflex gains of the leg extensor muscles during split-belt locomotion. **a**. H-reflexes and SLA adapt similarly when unilateral speed changes are introduced during split-belt locomotion (adapted from Refy et al., 2023). **b**. Schematic of H-reflex measurement with M- and H-wave responses highlighted (top), and definition of SLA (bottom).

Here we use a neuromuscular model of human walking on a split-belt treadmill to investigate the causal relationship between SLA and spinal reflex gain adaptation. The model’s control primarily consists of muscle reflexes. Using computer simulations, we adapt the gains of these reflexes in different combinations along with the belt speeds of the simulated treadmill. We find that in the model, reflex gain adaptation causes gait asymmetry rather than is a result of it. We also find that slow gain facilitation accounts for the reductions in gait asymmetry that occur during adaptation to the split belt condition, a feature absent in cerebellar degeneration patients (Morton and Bastian, 2006). Finally, we discuss the implications of these findings for the physiological processes underlying gait adaptation in the human motor system.

## 2 Methods

### 2.1 Neuromuscular Model

We used a 2-D version of a previously published neuromuscular model of human walking (Geyer and Herr, 2010; Song and Geyer, 2015) (Fig. 2-a). The model can walk at different speeds and stabilize against gait disturbances such as sudden speed changes of a simulated treadmill (Song and Geyer, 2017). The model includes the major leg muscle groups involved in sagittal plane walking. Its neural control distinguishes between stance and swing. During swing, the control drives the leg into a target pose that is adjusted based on stance width (Δ*x*) and center of mass speed (*ν* _*CM*_) to stabilize gait. During stance, the control relies on reflexes based on proprioceptive (muscle lengths and forces), vestibular (trunk lean), and mechano-receptor (perceived leg loading and foot contact) inputs to generate locomotion behavior. The strength of an individual reflex is characterized by a gain *G*^*i*^. For instance, the soleus muscle stimulation *s*(*t*) in the model is controlled by a time-delayed, positive force feedback, *s*_SOL_(*t*) = *s*_0_+*G*^SOL^ *F*^SOL^(*t*−Δ*t*), whose strength is determined by the reflex gain *G*^SOL^.

**Figure 2.**
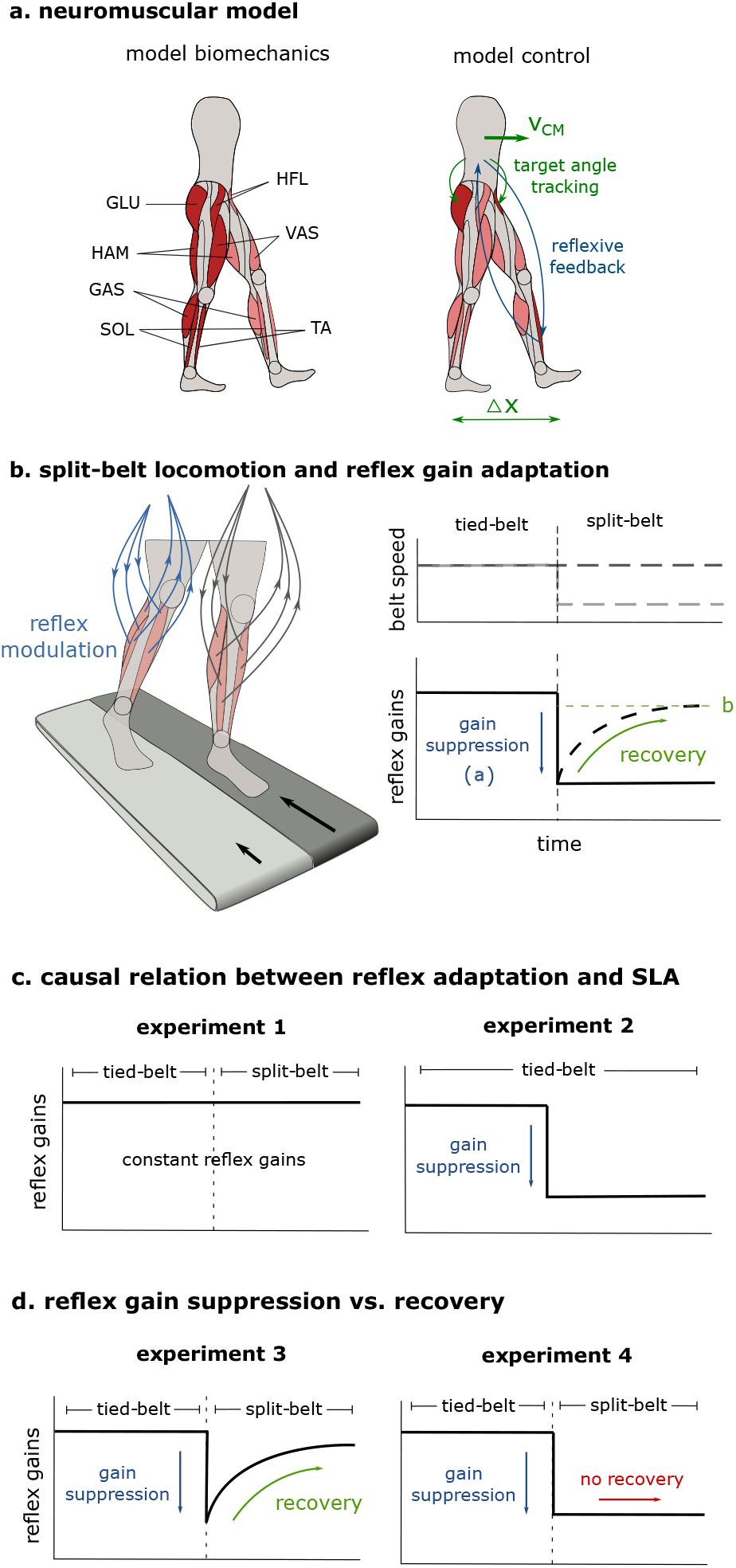
Overview of neuromuscular model and simulation experiments with four combinations of belt speed changes and reflex gain adaptations. **a**. 2-D neuromuscular model with leg muscles (GAS: gastrocnemius, SOL: soleus, TA: tibialis anterior, BFsH: biceps femoris short head, HAM: hamstrings, GLU: gluteus maximus, HFL: hip flexors) and main control elements indicated. **b**. Reflex gain adaptation function (Eq. 1). **c**. Simulation experiments 1 and 2 for studying causal relationship between gait asymmetry and reflex gain adaptation. **d**. Simulation experiments 3 and 4 for studying the roles of reflex gain suppression and recovery during split-belt locomotor adaptation.

### 2.2 Reflex Gain Adaptation

To explore the causal relationship between reflex gain adaptation and SLA, we introduced in the model an over-all gain adaptation 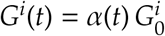 with

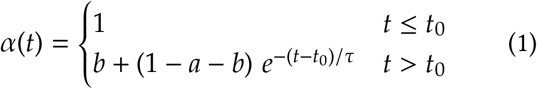

to all the reflexes of the leg on the belt undergoing speed changes (ipsilateral leg). The adaptation is parameterized by an initial gain suppression *a* at the time *t*_0_ of the belt speed change, a gain recovery time constant τ (τ = 10 s throughout the paper), and a steady-state level *b* at which the reflex gains settle after adaptation (Fig. 2-b). Although motivated by the time course of the H-reflex changes seen in the experiments (compare Fig. 1), α(*t*) can as well represent persistent suppression without recovery in the reflex gains (*b* = 1 − *a*) and no gain adaptation at all (*a* = 0, *b* = 1).

### 2.3 Spinal Gain Optimization

We used optimization of the initial gains 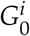 to identify walking solutions of the neuromuscular model when it is subjected to different experimental conditions (simulation experiments 1 through 4 indicated in Fig. 2c-d and detailed in Secs. 3.1 through 3.3). Specifically, we used simple Bayesian optimization with Gaussian process prior (Rasmussen and Williams, 2005) to minimize the cost function

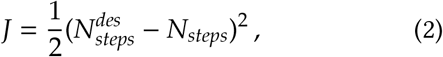

where 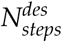 is a desired number of steps (set to 32 steps for a maximum simulated time of 40 seconds throughout this work), and *N*_*steps*_ denotes the actual steps taken by the model. At the start of each optimization, we set the gain parameters *G*^*i*^ to baseline values reported in earlier work (Geyer and Herr, 2010; Song and Geyer, 2015) and initialized the standard deviation of the Gaussian noise in the parameter space to 50% of these values. The optimization then repeatedly generated five new parameter sets by adding Gaussian noise to the baseline values, simulated the model for each parameter set, and chose the best performing set as the new baseline. With each iteration, we halved the standard deviation. In addition, we limited the range of the reflex gain values to between 0% and 250% of their initial values. For each experiment, we repeated the optimization five times to account for its stochastic nature and performed statistical analysis (Sec. 2.5) of this group of optimization runs when investigating SLA changes in different experimental conditions.

### 2.4 Definition of Step Length Asymmetry

We measured SLA in the model according to the conventions used in the literature (Morton and Bastian, 2006; Torres-Oviedo and Bastian, 2010). For two subsequent steps in the model, we used the distance between the left and right ankles at their respective heel strikes to define the step length *L*_*step*_. We then measured SLA as the difference between the left and right (or vice versa) step lengths, using

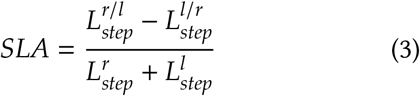

in simulations with tied belts of the treadmill (experiment 1) and

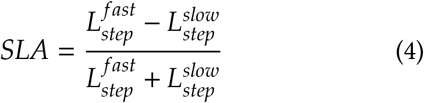

in simulations with split belts (experiments 2-4), where ‘fast’ and ‘slow’ refer to the leg moving on the faster and slower belt, respectively (Fig. 1).

### 2.5 Statistical Analysis

To compare the SLA adaptation observed in the four simulation experiments, we used several statistical tests. Specifically, we used two-way ANOVA (Ali and Bhaskar, 2016) to compare the same time intervals across the simulated experiments (Sec. 3.1) and one-way ANOVA to compare between different time intervals of each simulated experiment (Sec. 3.2). If ANOVA yielded a statistically significant difference in SLA, we performed post-hoc t-tests to identify statistically significant differences among different time intervals. Additionally, we corrected the p-values of t-post-hoc t-tests for false discovery rates using the Benjamini-Hochberg method (Benjamini and Hochberg, 1995) and used one-sample t-tests to check if the SLA is different from zero during tied-belt walking. Throughout this paper, we used 0.05 as the cut-off p-value for statistical significance.

## 3 Results

### 3.1 SLA effect of reflex gain adaptation rather than belt speed change

To better understand what causes SLA, we performed two simulation experiments. In the first experiment (experiment 1), we lowered the speed of the ipsilateral belt 10 seconds into the simulation by 30% (*t*_0_ = 10 s) but left the reflex gains of the model constant throughout (*a* = 0, *b* = 1). In the second experiment (experiment 2), we did not change the belt speed but instead lowered the reflex gains at *t*_0_ by 20% on the ipsilateral leg of the model (*a* = 0.2, *b* = 0.8). We expected either asymmetry to induce SLA in the model. However, the walking solutions that the neuromuscular model converged to express asymmetric gait only in response to reflex gain changes.

Figure 3 summarizes the SLA changes observed for the optimized neuromuscular model (mean and s.d. from five optimization runs per experiment) in the two experiments. Analyzing the SLA among four time intervals (right before and right after *t*_0_, midway through and at the end of the recovery phase), we find statistically significant interaction between the two groups of solutions (two-way ANOVA, *F* = 2.81, *p* = 0.002). We further observe that the model solutions in experiment 1 show increased SLA variability when introducing asymmetric belt speeds but no statistically significant trend (*p* > 0.05 between all intervals) (Fig. 3-a). In contrast, model solutions in experiment 2 consistently show significant changes in SLA when the reflex gains drop at *t*_0_ (post-hoc t-tests: *p* = 0.00016, *p* = 0.0018 and *p* = 0.0005 between the first and all subsequent intervals, respectively) but not thereafter (*p* > 0.25 between intervals 2-4) (Fig. 3-b). Thus, consistent trends in SLA are caused by changes in the reflex gains rather than by the mechanical asymmetry introduced with belt speed changes.

**Figure 3.**
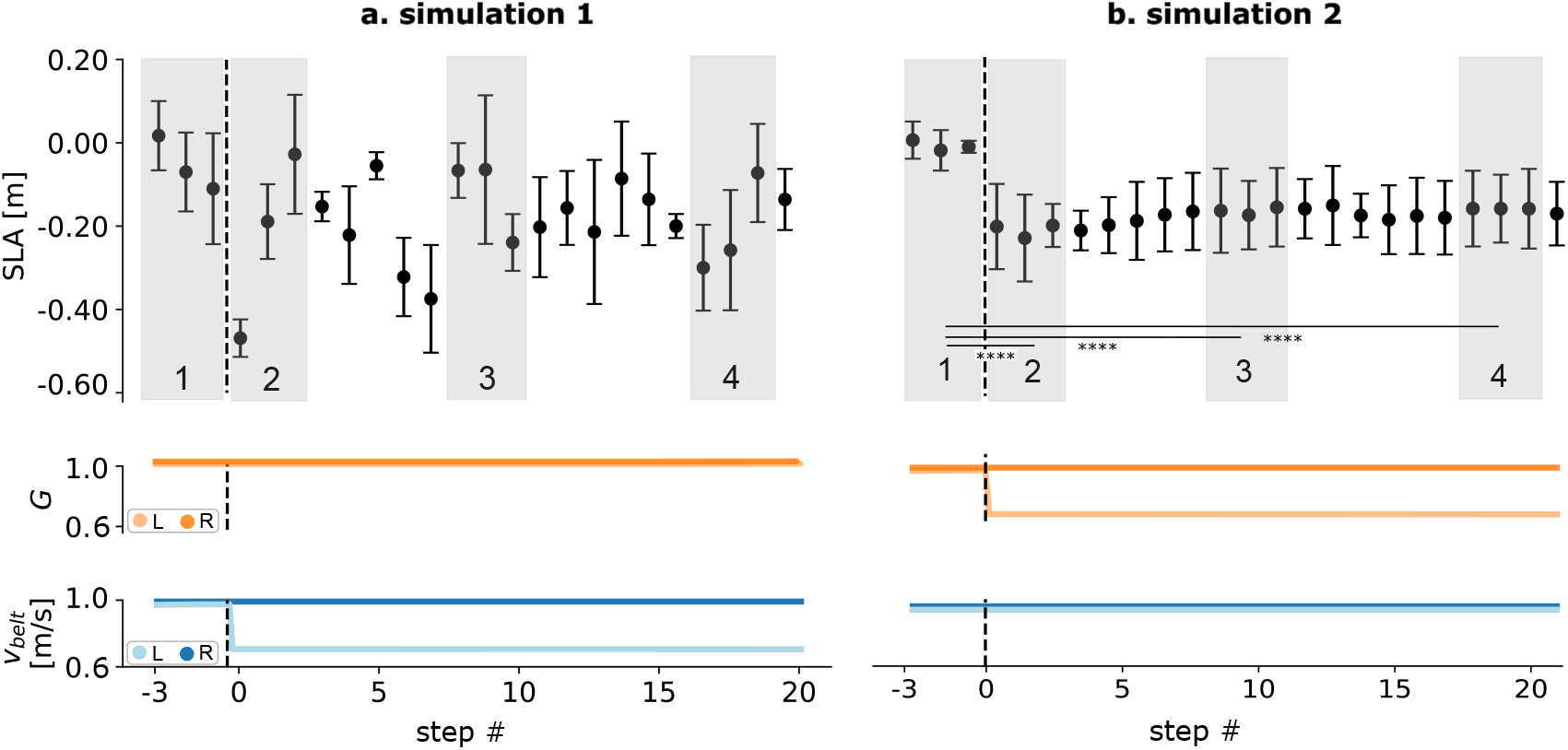
Cause of SLA changes. **a**. When imposing belt speed changes without reflex gain adaptation, model solutions show increased variability in SLA but no trend. **b**. When imposing reflex gain adaptation without belt speed changes, model solutions show clear change in SLA. Error bars represent one s.d.; ****: *p* < 0.0001.

### 3.2 Imposing gain suppression followed by recovery leads to SLA pattern observed in healthy adults

To further test if reflex adaptation could effect the SLA pattern observed in our previous human study (Refy et al., 2023) (Fig. 1), we optimized the neuromuscular model for walking with changes in belt speed and reflex gains similar to the ones observed in that study (experiment 3). Specifically, we optimized the model to walk on a simulated treadmill with an initially tied belt speed of 1.0 m s^−1^ that dropped by 30% on the ipsilateral side after 10 seconds. At the same time, the reflex gains of the model’s ipsilateral leg adapted according to (1) with *t*_0_ = 10 s, *a* = 0.30, and *b* = 0.90, mimicking the changes observed for the H-reflex in our human study (sudden gain drop by 80% at speed change followed by recovery to 90% of initial gain within about 45 seconds, compare Fig. 1).

The mean and standard deviation of the SLA patterns obtained from five optimization runs is summarized in figure 4-a. There are statistically significant differences among the four time intervals introduced in the last section (one-way ANOVA, *F* = 57.9; *p* = 1.4×10^−19^). These differences occur not only between before (interval 1) and after the speed change (intervals 2-4, post-hoc t-tests, *p* = 6.8 × 10^−8^, *p* = 3.2 × 10^−9^ and *p* = 1.5 × 10^−5^, respectively) but also between right after the speed change (interval 2) and later (intervals 3 and 4, *p* = 0.002 and *p* = 2.1 × 10^−5^, respectively), and between midway through the recover (interval 3) and at its end (interval 4, *p* = 2.6 × 10^−4^). Overall, the SLA patterns that the model solutions produce in repeated optimization runs are consistent and change with progress in recovery time. They also resemble the SLA pattern observed in our human study (Refy et al., 2023) (compare Fig. 1) as well as reported in the literature (Reisman et al., 2005; Morton and Bastian, 2006; Reisman et al., 2007; Torres-Oviedo and Bastian, 2010).

**Figure 4.**
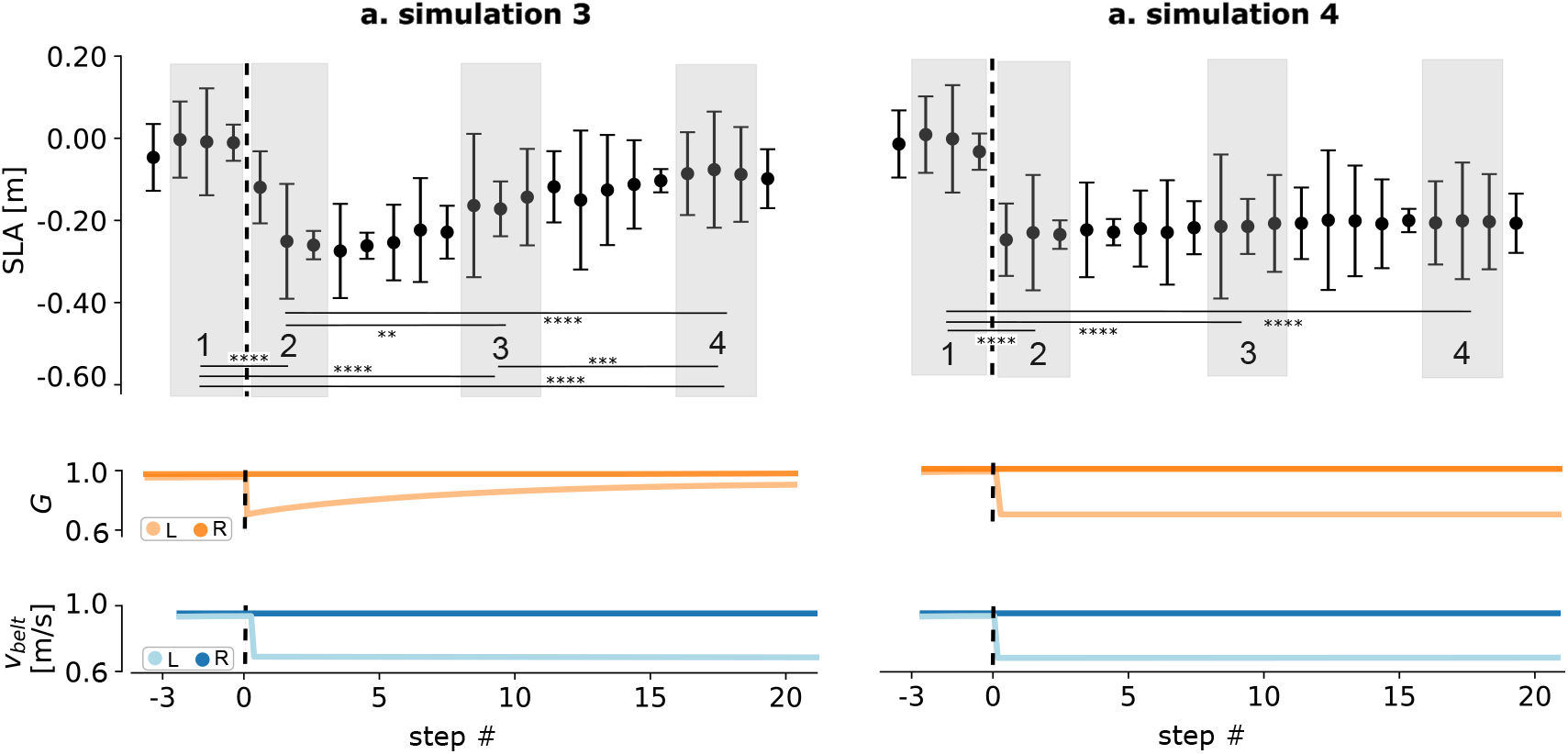
Reproducing SLA patterns observed in healthy young adults and cerebellar degeneration patients. **a**. With reflex gain adaptation similar to observed H-reflex changes, model solutions converge to SLA pattern of healthy young adults (compare Fig. 1). **b**. Without gain recovery after belt speed transition, model solutions show no long-term adaptation of SLA, a pattern observed in cerebellar degeneration patients. Error bars represent one s.d.; **: *p* < 0.01, ***: *p* < 0.001, ****: *p* < 0.0001.

### 3.3 Imposing gain suppression without recovery leads to SLA pattern observed in cerebellar degeneration patients

Cerebellar degeneration patients show a different pattern of SLA when they are exposed to ipsilateral belt speed reductions in treadmill locomotion experiments. Although their SLA rapidly changes when the speed reduces, it does not recover to baseline over time (Morton and Bastian, 2006). We found that we can reproduce this behavior in the model when we optimize it with the reflex gains unchanged after their initial drop (*a* = 0.25 and *b* = 0.75 in Eq. 1) (experiment 4, Fig. 4). In this case, the repeated optimization led to solutions whose SLA is significantly different before and after transition (one-way ANOVA: *F* = 50.6; *p* = 1.9×10^−16^; post-hoc t-tests: *p* = 4.2×10^−6^, *p* = 2.7 × 10^−6^, and *p* = 4.9 × 10^−6^, between intervals 1 and 2-4, respectively) but showed no significant difference between the intervals following the transition (*p* > 0.2 for all tests among intervals 2-4). Hence, as for cerebellar degeneration patients, the SLA did not recover to baseline.

## 4 Discussion

We used a neuromuscular model of human gait to study the causal relationship between spinal reflex adaptation and gait asymmetry in split-belt treadmill locomotion. Optimizing the model to walk in simulated treadmill experiments that isolate the effects of reflex adaptation and belt speed changes, we found that consistent trends in gait asymmetry emerge from the former and not the latter (Fig. 3). Moreover, we found that the model produces SLA patterns similar to healthy adults (Reisman et al., 2005, 2007; Torres-Oviedo and Bastian, 2010; Refy et al., 2023) when the reflex gains adapt in a way similar to the H-reflex changes we observed in a previous human study (Refy et al., 2023) (Fig. 4-a). This adaptation is characterized by a gain suppression immediately after the ipsilateral belt speed is reduced followed by a longer recovery toward the gain before the speed change. Finally, we demonstrated that if the recovery part of the adaptation is not taking place in the model, it produces SLA patterns similar to those observed for cerebellar degeneration patients (Morton and Bastian, 2006) (Fig. 4-b).

Our results support the hypothesis that spinal reflex adaptation *directly* causes gait asymmetry in human locomotion. Previous studies have reported an indirect causal link; more specifically, Thompson et al. (2013) found that down-conditioning the soleus H-reflex in spinal cord injured people can reduce gait asymmetry, corroborating earlier observations made by Chen et al. (2006) in spinal cord injured rats. However, these changes in gait asymmetry occurred over the course of months and were likely the outcome of a reduced hyper-reflexia due to the down-conditioning (Thompson and Wolpaw, 2014, 2021). Our recent observation of near synchronous changes in the soleus H-reflex and the SLA during split-belt locomotion indicated a closer link between spinal reflex adaptation and gait asymmetry (Refy et al., 2023) (Fig. 1), and the modeling results obtained here provide evidence for changes in gait asymmetry to be directly caused by reflex adaptation.

The functional purpose of reflex adaptation and what mechanisms regulate it remain unclear. In the neuromuscular model, the adaptation was imposed by us (Eq. 1), and without it, the model optimization produced walking solutions that were less stable. More exactly, these solutions had walking patterns with somewhat erratic large and small steps due to repeated near stumbles, which accounts for the enlarged SLA variability observed in the first simulation experiment (Fig. 3-a). Thus, in the model, reflex adaptation is needed to maintain stable walking, especially, right after the belt speed change. Whether humans adapt their reflexes for the same reason remains unclear. Although the predictive capabilities of the neuromuscular model (Song and Geyer, 2017, 2018) and similar other ones (Dorn et al., 2015; Ong et al., 2019) have been demonstrated to some extent, these models clearly simplify the human motor control system. For instance, the human system could include oligosynaptic spinal circuits based on the stretch velocity of the hip muscles that modulate the gains of simpler reflex circuits in proportion to the current walking speed (Meunier et al., 1996; Misiaszek and Pearson, 1997). Such a layered network would necessarily change the gains of the simpler reflexes at the belt speed change, and the observed gain recovery thereafter could reflect a drift toward a new gain equilibrium as the gait biomechanics adjust to the new situation. In this scenario, none of the gain adaptation would be in response to a less stable walk.

Some indirect evidence that the immediate gain change at the belt speed transition may indeed be of spinal origin comes from animal studies. Cats respond to belt speed changes in split-belt locomotion with an asymmetric gait, although they lack the longer term recovery toward gait symmetry seen in humans. This adaptation pattern is likely of spinal origin, as it persists in spinalized cats (Kuczynski et al., 2017). Furthermore, the pattern also shows for the modulation of cutaneous reflexes, which are reduced in correlation with the belt speed asymmetry in both the intact and spinalized cat Hurteau and Frigon (2018). Together, these observations are consistent with the idea that the change of the reflex gains, and hence the gait asymmetry, at the belt speed transition is an immediate reaction of the spinal circuitry.

On the other hand, our observation that the model’s SLA pattern matches those of cerebellar degeneration patients (Morton and Bastian, 2006) when its gain recovery is removed (Fig. 4-b) suggests that the gain recovery is regulated by supraspinal rather than spinal inputs. The cerebellum plays a critical role in modulating reflexes in other scenarios; for instance, it has been shown that cerebellar lesions can prevent the ability of rats to down-condition their lower limb H-reflexes (Chen and Wolpaw, 2005) as well as that such lesions eliminate conditioned eye blink reflexes in rabbits and saccadic eye movements in monkeys (McCormick and Thompson, 1984; Barash et al., 1999). Closer to our current results, MacKay and Murphy (1979) reviews substantial evidence that the cerebellum continuously modulates the gains of proprioceptive reflexes involved in limb movement. Given our model observations, we speculate that the cerebellum also regulates the reflex gain recovery in human split belt walking, and that the H-reflexes in patients with cerebellar degeneration will suddenly change at the belt speed change but not recover thereafter.

In summary, the overall picture that emerges is that the dynamic adaptation of spinal reflex gains may be an essential part of human gait control when encountering unexpected external changes such as the speed changes during split-belt locomotion, and that this adaptation has two components, an immediate response perhaps originating from the spinal circuitry itself and a longer-term recovery governed by the cerebellum (Fig. 5).

**Figure 5.**
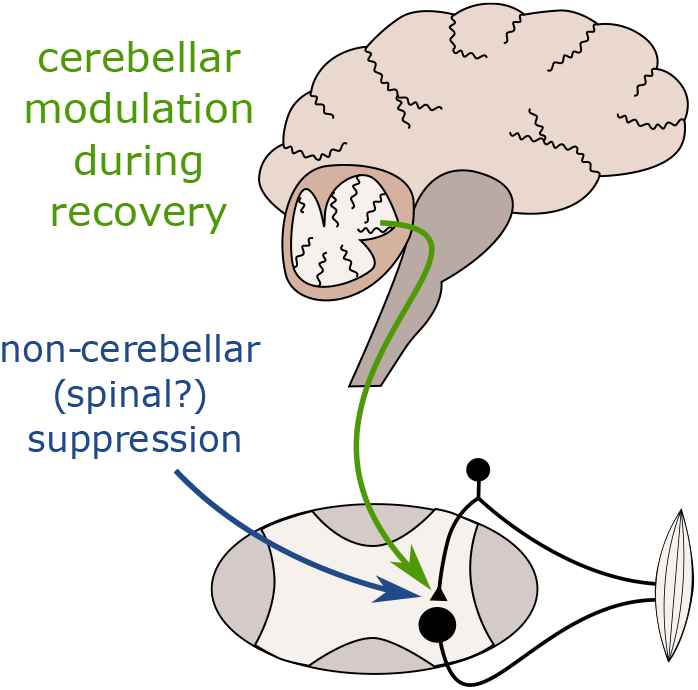
Overall picture of spinal gain adaptation during split-belt locomotion. Our results suggest the cerebellum modulates spinal reflex gains during the SLA recovery phase, while the initial gain suppression is of non-cerebellar origin.

## 5 Acknowledgements

The authors would like to acknowledge Meadow Webster for drawing most illustrations of this paper.

## References

Ali, Z. and Bhaskar, S. B. (2016). Basic statistical tools in research and data analysis. Indian J. Anaesth., 60(9):662–669.

Barash, S., Melikyan, A., Sivakov, A., Zhang, M., Glickstein, M., and Thier, P. (1999). Saccadic dysmetria and adaptation after lesions of the cerebellar cortex. The Journal of Neuroscience, 19:10931–10939.

Benjamini, Y. and Hochberg, Y. (1995). Controlling the false discovery rate: A practical and powerful approach to multiple testing. Journal of the Royal Statistical Society. Series B (Methodological), 57(1):289–300.

Chen, X. Y. and Wolpaw, J. R. (2005). Ablation of cerebellar nuclei prevents h-reflex down-conditioning in rats. Learning & Memory, 12:248–254.

Chen, Y., Chen, X. Y., Jakeman, L. B., Chen, L., Stokes, B. T., and Wolpaw, J. R. (2006). Operant conditioning of h-reflex can correct a locomotor abnormality after spinal cord injury in rats. Journal of Neuroscience, 26:12537–12543.

di Biase, L., Di Santo, A., Caminiti, M. L., De Liso, A., Shah, S. A., Ricci, L., and Di Lazzaro, V. (2020). Gait Analysis in Parkinson’s Disease: An Overview of the Most Accurate Markers for Diagnosis and Symptoms Monitoring. Sensors, 20(12):3529.

Dietz, V., Zijlstra, W., and Duysens, J. (1994). Human neuronal interlimb coordination during split-belt locomotion. Experimental Brain Research, 101(3):513–520.

Dorn, T. W., Wang, J. M., Hicks, J. L., and Delp, S. L. (2015). Predictive Simulation Generates Human Adaptations during Loaded and Inclined Walking. PLOS ONE, 10(4):e0121407.

Geyer, H. and Herr, H. (2010). A muscle-reflex model that encodes principles of legged mechanics produces human walking dynamics and muscle activities. IEEE Transactions on Neural Systems and Rehabilitation Engineering, 18:263–273.

Hurteau, M.-F. and Frigon, A. (2018). A spinal mechanism related to left–right symmetry reduces cutaneous reflex modulation independently of speed during split-belt locomotion. The Journal of Neuroscience, 38:10314–10328.

Iturralde, P. A. and Torres-Oviedo, G. (2019). Corrective muscle activity reveals subject-specific sensorimotor recalibration. eNeuro, 6(2).

Jacobsen, N. A. and Ferris, D. P. (2023). Electrocortical activity correlated with locomotor adaptation during split-belt treadmill walking. The Journal of Physiology, 601(17):3921–3944.

Kuczynski, V., Telonio, A., Thibaudier, Y., Hurteau, M., Dambreville, C., Desrochers, E., Doelman, A., Ross, D., and Frigon, A. (2017). Lack of adaptation during prolonged split-belt locomotion in the intact and spinal cat. The Journal of Physiology, 595:5987–6006.

Li, S., Francisco, G. E., and Zhou, P. (2018). Poststroke Hemiplegic Gait: New Perspective and Insights. Frontiers in Physiology, 9.

MacKay, W. and Murphy, J. (1979). Cerebellar modulation of reflex gain. Progress in Neurobiology, 13(4):361–417.

McCormick, D. A. and Thompson, R. F. (1984). Cerebellum: Essential involvement in the classically conditioned eyelid response. Science, 223:296–299.

Meunier, S., Mogyoros, I., Kiernan, M. C., and Burke, D. (1996). Effects of femoral nerve stimulation on the electromyogram and reflex excitability of tibialis anterior and soleus. Muscle & Nerve, 19(9):1110–1115.

Misiaszek, J. E. and Pearson, K. G. (1997). Stretch of quadriceps inhibits the soleus h reflex during locomotion in decerebrate cats. Journal of Neurophysiology, 78(6):2975–2984. PMID: 9405517.

Morton, S. M. and Bastian, A. J. (2006). Cerebellar contributions to locomotor adaptations during splitbelt treadmill walking. Journal of Neuroscience, 26.

Nas, K. (2015). Rehabilitation of spinal cord injuries. World Journal of Orthopedics, 6(1):8.

Ong, C. F., Geijtenbeek, T., Hicks, J. L., and Delp, S. L. (2019). Predicting gait adaptations due to ankle plantarflexor muscle weakness and contracture using physics-based musculoskeletal simulations. PLOS Computational Biology, 15(10):e1006993.

Rasmussen, C. E. and Williams, C. K. I. (2005). Gaussian Processes for Machine Learning. The MIT Press.

Refy, O., Blanchard, B., Miller-Peterson, A., Dalrymple, A. N., Bedoy, E. H., Zaripova, A., Motaghedi, N., Mo, O., Panthangi, S., Reinhart, A., Torres-Oviedo, G., Geyer, H., and Weber, D. J. (2023). Dynamic spinal reflex adaptation during locomotor adaptation. Journal of Neurophysiology, 130(4):1008–1014. PMID: 37701940.

Reisman, D. S., Block, H. J., and Bastian, A. J. (2005). Interlimb coordination during locomotion: What can be adapted and stored? Journal of Neurophysiology, 94.

Reisman, D. S., Wityk, R., Silver, K., and Bastian, A. J. (2007). Locomotor adaptation on a split-belt treadmill can improve walking symmetry post-stroke. Brain, 130:1861–1872.

Song, S. and Geyer, H. (2015). A neural circuitry that emphasizes spinal feedback generates diverse behaviours of human locomotion. The Journal of Physiology, 593(16):3493–3511.

Song, S. and Geyer, H. (2017). Evaluation of a neurome-chanical walking control model using disturbance experiments. Frontiers in Computational Neuroscience, 11.

Song, S. and Geyer, H. (2018). Predictive neuromechanical simulations indicate why walking performance declines with ageing. The Journal of Physiology, 596(7):1199–1210.

Thompson, A. K., Pomerantz, F. R., and Wolpaw, J. R. (2013). Operant Conditioning of a Spinal Reflex Can Improve Locomotion after Spinal Cord Injury in Humans. The Journal of Neuroscience, 33(6):2365–2375.

Thompson, A. K. and Wolpaw, J. R. (2014). Operant conditioning of spinal reflexes: from basic science to clinical therapy. Frontiers in Integrative Neuroscience, 8.

Thompson, A. K. and Wolpaw, J. R. (2021). H-reflex conditioning during locomotion in people with spinal cord injury. The Journal of Physiology, 599:2453–2469.

Torres-Oviedo, G. and Bastian, A. J. (2010). Seeing is believing: Effects of visual contextual cues on learning and transfer of locomotor adaptation. Journal of Neuroscience, 30:17015–17022.

Vazquez, A., Statton, M. A., Busgang, S. A., and Bastian, A. J. (2015). Split-belt walking adaptation recalibrates sensorimotor estimates of leg speed but not position or force. Journal of Neurophysiology, 114(6):3255–3267. PMID: 26424576.

